# Relationship between advanced footwear technology longitudinal bending stiffness and energy cost of running

**DOI:** 10.1101/2023.12.27.573423

**Authors:** Víctor Rodrigo-Carranza, Wouter Hoogkamer, José María González-Ravé, Fernando González-Mohíno

**Affiliations:** Sports Performance Research Group (GIRD) University of Castilla-La Mancha, Toledo, Spain; Integrative Locomotion Laboratory, Department of Kinesiology, University of Massachusetts, Amherst, MA, USA; Facultad de Ciencias de la Vida y de la Naturaleza, Universidad Nebrija, Madrid, Spain

**Keywords:** running shoes, running economy, performance, rocker, super shoes

## Abstract

**Introduction/Purpose:** Shoe longitudinal bending stiffness (LBS) is often considered to influence running economy (RE) and thus, running performance. However, previous results are mixed and LBS levels have not been studied in advanced footwear technology (AFT). The purpose of this study was to evaluate the isolated effects of increased LBS from curved carbon fiber plates embedded within an AFT midsole on RE and spatiotemporal parameters.

**Methods:** Twenty-one male trained runners completed 3 times 4 minutes at 13km/h with two experimental shoe models with a curved carbon fiber plate embedded in an AFT midsole with different LBS values (Stiff: 35.5 N/mm and Stiffest: 43.1 N/mm), and a Control condition (no carbon fiber plate: 20.1 N/mm). We measured energy cost of running (W/kg) and spatiotemporal parameters in one visit.

**Results:** RE improved for the Stiff shoe condition (15.71 ± 0.95 W/kg; p<0.001; *n*^*2*^ =0.374) compared to the Control condition (16.13 ± 1.08 W/kg) and Stiffest condition (16.03 ± 1.19 W/kg). However, we found no significant differences between the Stiffest and Control conditions. Moreover, there were no spatiotemporal differences between shoe conditions.

**Conclusion:** Changes in LBS in AFT influences RE suggesting that moderately stiff shoes are the most effective LBS to improve RE in AFT compared to very stiff shoes and traditional, flexible shoe conditions while running at 13 km/h.

## INTRODUCTION

Running performance, particularly in long distance events such as the marathon, depends on an interplay of several physiological factors ^1,2^, including running economy (RE). The metabolic cost of running is dominated by the cost of generating force to support body weight ^3^ and theoretically can be improved by minimizing the mechanical energy loss or absorption of muscles and tendons to lift and accelerate the body and limbs ^3^. Advanced footwear technology (AFT) shoes are characterized by the presence of a curved carbon fiber plate [with the aim of optimizing longitudinal bending stiffness (LBS; greater resistance to bending of the shoe) to minimize mechanical energy losses at the metatarsophalangeal joints ^4,5^] embedded within a resilient, compliant and lightweight midsole foam (e.g., PEBA) to provide cushioning and maximize mechanical energy return ^6^. The combination of these features has demonstrated benefits for distance running performance ^7–10^ through RE improvements ^6,11^. The RE improvements from AFT shoes are usually accompanied by small spatiotemporal changes (i.e., an increase in step length and contact time) ^4,5^.

Previous studies on the effects of increased LBS on RE found that the first AFT shoes (the Nike Vaporfly and its preceding prototype) improved RE between 2 and 6% compared to traditional racing shoes^6,11^. To study the *isolated* effects of increased LBS on the metabolic cost of running (RE), several studies have used baseline control shoes and modified LBS by embedding carbon-fiber plates in the midsole or using carbon-fiber insoles, keeping other properties (e.g., upper, midsole foam and geometry) constant. There appears to be a distinction between studies that used embedded carbon-fiber plates and reported significant improvements in RE of around 1% ^12–14^ versus studies that use top-loaded plates (insoles) and did not find overall significant RE improvements ^15–18^. Roy & Stefanyshyn ^14^ studied traditional running shoes with three different levels of LBS using flat carbon fiber plates embedded in the midsole for running on a treadmill at ∼13.5 km/h. They found that compared to control shoes (18 N/mm), the oxygen cost decreased by 0.8 % when wearing shoes with a stiff (38 N/mm) carbon fiber plate. In contrast, the oxygen cost when wearing shoes with even stiffer carbon fiber plates (45 N/mm) was not significantly different compared to control shoes ^14^. Other studies have found no RE differences when increasing LBS using flat carbon fiber plates or insoles, even when controlling shoe mass and with LBS values similar to Nike Vaporfly shoes ^16–18^. The main difference of the shoe conditions in these studies compared to AFT shoes is that AFT shoes have a curved plate instead of a flat plate and more resilient and compliant foam. Farina et al., ^19^ showed that flat plates reduce negative metatarsophalangeal joint work, but increase ankle push-off moment demands, while curved plates reduce negative metatarsophalangeal joint work without increased ankle push-off demands. Altogether, this suggests that plate location and shape are important factors when LBS is increased ^4,19^. Healey and Hoogkamer ^20^ where the first to specifically investigate the effect of LBS in AFT (Nike Vaporfly). They did not compare plate vs. no plate, or plates of different levels of LBS, but made several medio-lateral cuts in the midsole through the carbon fiber plate of the AFT to reduce LBS. They observed a small not significant changes in energy cost. The combination of these results suggests that the increase in LBS cannot explain all the improvement running in AFT and the new midsole foams have an important role to play.

To further explore whether an optimal LBS exists in AFT, we studied the isolated effects of two levels of LBS in AFT versus a traditional, flexible shoe. Importantly, we used curved plates, embedded within a modern midsole foam, specific to AFT, the most commonly used type of shoe in major road events today ^21^. The purpose of this study was to evaluate the isolated effects of increased LBS by embedding a curved carbon fiber plate within the AFT midsole on RE and spatiotemporal parameters. Based on previous literature ^5,14^, we hypothesized that there would be differences in RE depending on the increase of LBS where intermediate stiffnesses would obtain the best result on RE. We anticipated that this will be accompanied by small changes in spatiotemporal variables such as an increase in contact time and a reduction in step frequency.

## MATERIALS AND METHODS

### Participants

A power analysis was performed a priori (G^*^Power 3.1; Universität Kiel, Kiel, Germany) based on RE results of a previous study ^22^. This analysis indicated that 20 runners were required to detect a moderate effect size (0.30) between conditions, with a β⍰= ⍰0.20 and α⍰= ⍰0.05. Twenty-one male runners (mean ± SD: 27.6 ± 5.2 years; 65.3 ± 5.7 kg and 175.3 ±4.7 cm) who were experienced AFT users participated in this study. The inclusion criteria were the following: 1) participation in endurance training for at least 3 days per week during the previous 6 months without injury, 2) wearing a men’s US 9 shoe size. Exclusion criterion was inability to run at 13 km/h while blood lactate accumulation was below 4 mmol/L and with respiratory exchange ratio (RER) values below 1.0 during all the tests.

Prior to the study, all participants were informed about the testing protocols, possible risks involved and were invited to provide written informed consent. The study was performed in accordance with the principles of the Declaration of Helsinki (October 2008, Seoul), and the experimental protocols were approved by the local ethics committee (CEIC924).

### Shoe characteristics

Two experimental shoe models with curved carbon fiber plates with different levels of LBS, with identical uppers, foam midsole hardness and rubber outsoles were manufactured for this study and compared to a different Control condition shoe model with a similar midsole without a carbon fiber plate. The senior researcher labelled the shoes with a red, blue or without mark according to the two experimental conditions (Stiff and Stiffest) and Control condition, respectively. This was done to keep the study participants blinded. Only the senior researcher and first author knew which condition each mark corresponded to, so that when collecting data, the other researchers, and runners themselves did not know which condition was tested. Shoe characteristics are presented in Table 1. The LBS of the shoes was determined by a 3-point bending test on the complete shoe (including shoe upper; figure 1) ^13,14^ where the forefoot of the shoe was placed on a custom-made structure with two supporting pins, 80 mm apart, to a material testing machine that measures force and displacement (Microtest, S.A., Madrid, Spain). The machine was set to displace the center point of the forefoot 20 mm vertically downward at a speed of 4mm/s, at which point it returned to its original position in the same amount of time. This was repeated ten times for each shoe condition. The slope of the force–displacement curve (i.e., stiffness) was determined for all ten loading cycles. The stiffness values were first averaged between 80 and 90% of each loading curve (i.e., linear portion of the force–displacement curve) and then across all ten cycles ^13,14^. The LBS was 20.1 N/mm for the Control, 35.5 N/mm for the Stiff and 43.1 N/mm for the Stiffest condition, similar to values previously used ^14,18^. The mass of the shoes was 229 g, 222 g and 221 g for Control and experimental conditions, respectively ^23^. In addition, the experimental shoes models were identical in the composition and characteristics (material, color, and shape), with the aim to avoid any possible placebo effects ^11^. The only exception was the LBS of the carbon fiber plate inserted in the experimental shoe models. Furthermore, the Control condition was a different model, but with similar features (Table 1). This model was selected because it has the same drop and midsole height as the experimental conditions but does not have a carbon fiber plate to increase the LBS. Previous studies have shown that these characteristics can influence the RE ^24,25^.

**Figure 1.**
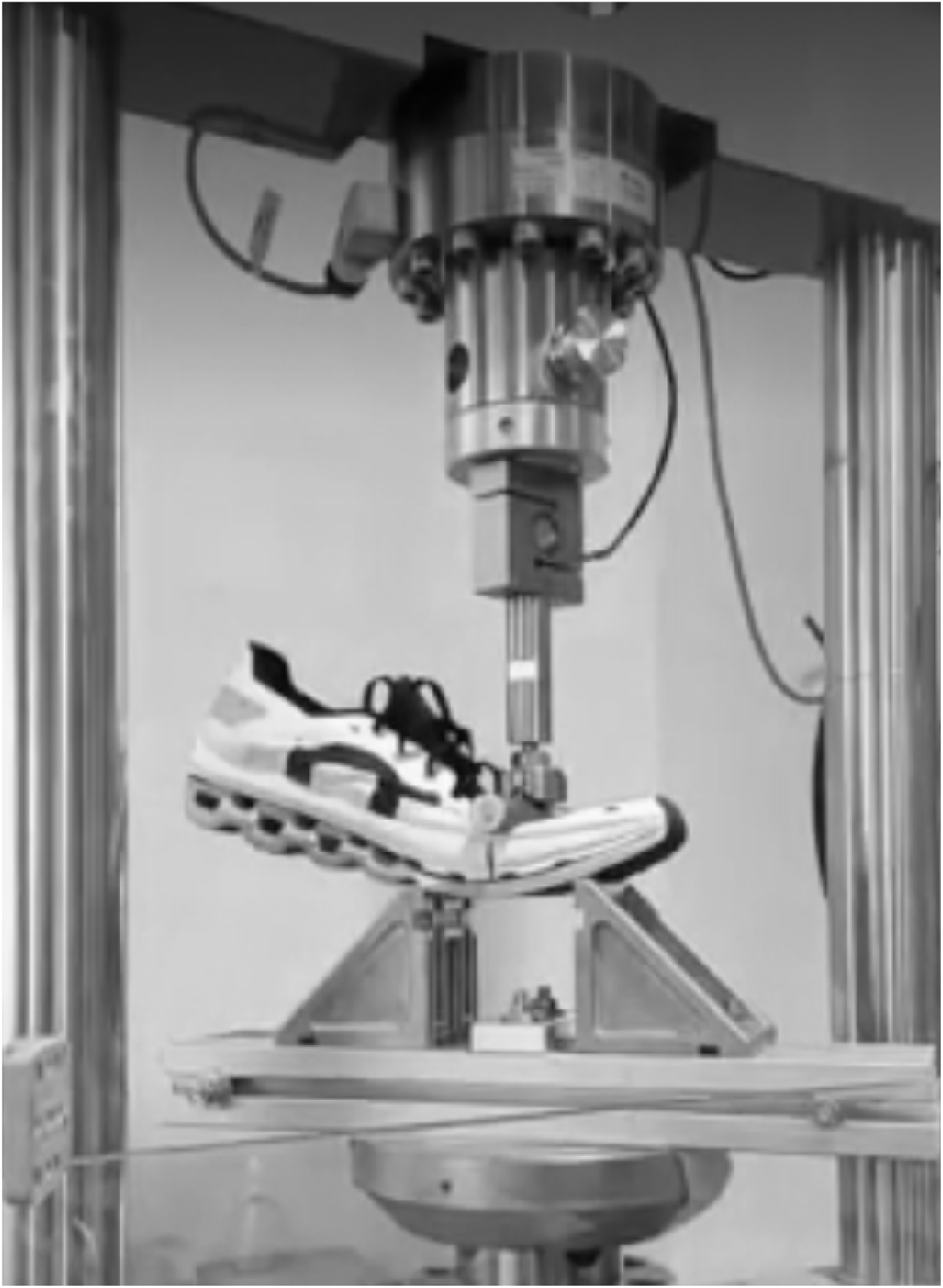
Set up of mechanical stiffness test on running shoes.

**Table 1.** Shoe characteristics and comfort of experimental conditions of size US 9.

| Shoe | Mass (g) | Carbon Plate | Heel stack (mm) | Forefoot stack (mm) | Drop (mm) | LBS (N/mm) |
| --- | --- | --- | --- | --- | --- | --- |
| Control | 229 | No | 38 | 29 | 9 | 20.1 |
| Stiff condition | 222 | Yes | 38 | 29 | 9 | 35.5 |
| Stiffest condition | 221 | Yes | 38 | 29 | 9 | 43.1 |

LBS, Longitudinal bending stiffness of the shoe

### Design and methodology

The effect of increased LBS on RE and spatiotemporal parameters was evaluated using a randomized crossover experimental design. All participants refrained from intense exercise in the 48h preceding the laboratory session (controlled environmental conditions [550m altitude, 20-22ºC, 35-40% relative humidity]). Participants were asked to avoid caffeine and alcohol intake within 24 hours of their testing visits and refrained from eating and drinking for 4h before testing.

First, anthropometrical variables were measured. Height was measured to the nearest 0.1 cm with a portable stadiometer (Seca, Bonn, Germany) and body mass was measured to the nearest 0.1 kg with a portable balance (Seca, Bonn, Germany), while barefoot and wearing lightweight shorts. Then, the researchers fitted the participant with the shoe size of all conditions, while the participant was not allowed to manipulate the footwear at any time. Prior to beginning the RE testing trials, all participants ran 10 min as a warm-up in their own shoes at a self-selected pace slower than 13 km/h. Participants then completed a 4 min trial at 13km/h with each shoe condition. This speed was selected such that participants were able to run below the lactate threshold to ensure steady-state O_2_ measurements and was similar to that used in previous studies that evaluated the optimal LBS with flat plates in traditional shoes ^14,17^. This was confirmed by blood lactate measures (values below 4 mmol/L) and RER (values below 1.0). All running trials were performed on a motorized treadmill (HP Cosmos Pulsar, HP Cosmos Sports & Medical GMBH, Nussdorf-Traunstein, Germany) with a vertical stiffness value of around 300 kN/m ^26^. The sequence of the shoes was randomized for each participant using a random number generator. During the RE trials, respiratory variables were measured using a gas analyzer (CPX Ultima Series MedGraphics, St. Paul, Minnesota, USA), which was calibrated prior to each session (CO_2_ 4.10%; O_2_ 15.92%). 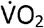 values collected during the last 60s of each 4 min bout were used to determine RE as oxygen cost of running ^27^. The energy cost of running was calculated (W/kg) from 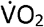 and 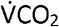 values ^28,29^. Blood samples were collected from the fingertip in the final minute before finishing the stage for blood lactate concentration determination (Lactate Scout, SensLab GmbH, Germany). After each trial, participants rested 10 minutes before the next trial.

The main spatiotemporal parameters of the gait cycle (contact time and step frequency) were measured for every step during treadmill running using a Stryd^®^ Power Meter device (V2, Stryd^®^, Boulder, CO, USA), sampling at 1000 Hz ^30^ and the information was analyzed using the web-based Stryd Power Center application. Data were analyzed for the final minute of each trial and was averaged for subsequent analyses. The 1-minute period provided well over the 25 steps recommended to distinguish running technique differences between people ^31^.

### Statistical analysis

Statistical analyses were carried out using the software IBM SPSS Statistics for Macintosh, Version 28.0 (IBM Corp., Armonk, NY, US). Data were screened for normality of distribution and homogeneity of variance using a Shapiro-Wilk Normality Test. Repeated-measures ANOVA was conducted to compare shoe conditions. When a significant main effect for shoe was observed, a Bonferroni *post hoc* test was performed. Moreover, repeated-measures ANOVA was conducted to compare the randomized effect of the distribution of the conditions. The effect size was calculated using the partial eta squared (ŋ^2^) in the repeated-measures ANOVA. Values of 0.01, 0.06 and above 0.15 were considered as small, medium and large, respectively ^32^. Significance level for all analyses was set at α = 0.05.

## RESULTS

### Metabolic measurements

All participants ran all trials with a lactate concentration below 4 mmol/L and it was similar between conditions (1.60 ± 0.63 mmol/L, 1.73 ± 0.69 mmol/L, 1.92 ± 1.19 mmol/L for Control, Stiff and Stiffest respectively; p=0.209; η^2^=0.079). RER remained below 1.0 across all trials for all participants (0.85 ± 0.08, 0.84 ± 0.08, 0.84 ± 0.09 for Control, Stiff and Stiffest respectively; p=0.21; η^2^=0.13). ANOVA revealed no significant differences (p = 0.992) in oxygen uptake 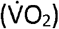 based on the testing sequence of the shoes, which indicates that the randomization assignment was effective.

There were significant differences between shoe conditions in RE measured as energy cost (p<0.001; η^2^ *= 0.374)*. RE improved for the Stiff shoe condition (15.71 ± 0.95 W/kg) compared to the Control condition (16.13 ± 1.08 W/kg; p <0.001) and Stiffest condition (16.03 ± 1.19 W/kg; p = 0.038). However, there were no differences between Stiffest and Control conditions (p = 0.241) (Figure 2).

**Figure 2.**
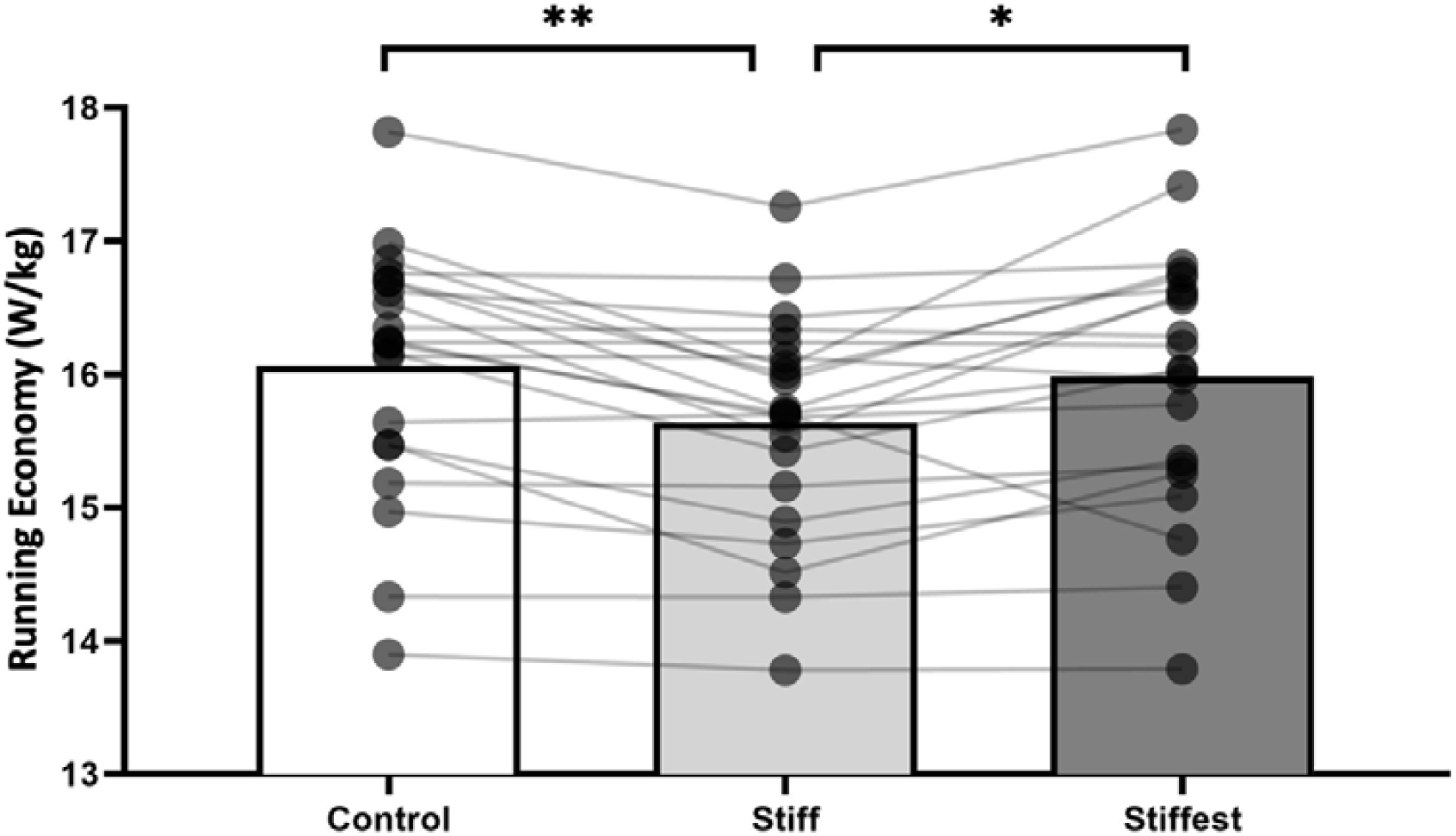
Running economy (W/kg). Bar graphs represent mean values and circles represent runners. ^*^p ≤ 0.05 and ^**^p ≤ 0.01 during post hoc comparisons when main effect of footwear significant. n =21

### Spatiotemporal measurements

No significant difference was detected in step frequency (167.2 ± 6.9; 166.7 ± 6.9; 166.9 ± 7.6 steps/min for Control, Stiff and Stiffest respectively; p=0.345; η^2^=0.054) and contact time between conditions (0.231 ± 0.014; 0.234 ± 0.010; 0.233 ± 0.010 s for Control, Stiff and Stiffest respectively; p=0.085; η^2^=0.121).

## DISCUSSION

The purpose of this study was to evaluate the isolated effects of increased LBS from curved carbon fiber plates with different LBS values, embedded within the AFT midsole on RE and spatiotemporal parameters. In support of our hypothesis, we found a significant decrease in energy cost of running (better RE) of 2.56% in the Stiff condition compared to Control and a significant decrease of 1.98% in the Stiff condition compared to the Stiffest condition. However, participants no experienced spatiotemporal changes when running with any of the shoe conditions. The relationship between the energy cost of running and the increase in LBS of AFT shoes we observed was similar (figure 2) but of greater magnitude (2.56% vs. 0.80%) in the Stiff condition over the Control condition to those previously reported by Roy and Stefanyshyn, ^14^ who used flat carbon fiber plates in traditional running shoes. This relationship suggests that an optimal LBS exists to improve RE, indicating that stiff AFT shoes (35.5 N/mm) are better than very stiff AFT shoes (43.1 N/mm), and than shoes without a carbon fiber plate (20.1 N/mm), to improve RE.

Our findings support the concept that carbon fiber plates are more effective in improving RE when plates are curved and embedded within a modern midsole foam ^4,19^. Shoes with an appropriate combination of LBS and plate curvature can reduce net mechanical energy loss at the metatarsophalangeal joint without increasing mechanical demands at the ankle ^4,19^. A curved plate has a higher effective LBS since it acts further from the point of rotation, creating a larger resultant moment around the metatarsophalangeal joints [see Fig. 1 ^4^]. In contrast, adding a flat carbon fiber plate to increase the LBS of the shoe typically shifts the runner’s center of pressure more anterior along the foot during ground contact ^33^. These altered biomechanics generally yield a longer moment arm between the ground reaction force and the ankle-joint center that leads to a larger ground reaction force-induced ankle-joint moment ^15^. Similarly, increasing LBS excessively could restrict the metatarsophalangeal joint’s natural dorsiflexion too much and would generate a longer moment arm to the ankle, which may explain the increase in energy cost (∼2%) in the Stiffest condition compared to the Stiff condition ^15,18^.

The improvements in RE found in this study between Stiff and Control conditions was bigger than in Roy and Stefanyshyn ^14^ in spite of using shoe conditions of similar LBS and at the same speed. Our results are more comparable with the results of running in AFT (compliant midsole with curved plates) compared to traditional running shoes with an improvement in RE of around 3-4%. ^6,11^. There is a growing body of literature indicating that the new more resilient and compliant midsole foams are of greater importance than increased LBS in improving RE when using AFT ^20,34,35^. Recently, we showed on separate occasions that AFT shoes with a curved carbon fiber plate and a highly resilient and compliant midsole (PEBA) improved RE by 1-2% over shoes with curved carbon fiber plate and traditional midsole (EVA) ^36,37^. It may be that the ∼2.5% improvement from the Stiff versus Control condition in the current study resulted from a ∼1% improvement from increased LBS (similar to reported in previous studies ^14,15^) and the remainder of the improvement from using more compliant and resilient midsole materials in combination with the curved carbon fiber plate. Therefore, it appears the combination of these characteristics of AFT is key to improve RE ^4,35^.

Regarding spatiotemporal parameters, the Stiff and Stiffest condition showed no significant differences with respect to the Control condition. However, there was a non-significant increase (p=0.085; η^2^=0.121) in contact time: the Stiff condition increased 1.30% and the Stiffest condition 0.86% compared to the Control condition. In a meta-analysis, we showed that improvements in RE from increasing LBS are accompanied by increases in step length and contact time ^5^. By increasing the contact time, the lower limb muscles have more time to contract in a stiff shoe and could therefore shorten at slower velocities compared with a control shoe ^38,39^, which is related to better movement economy. A more detailed biomechanical analysis of joint mechanics ^20,40^ could provide more insights into the mechanisms of how different levels of LBS affects RE.

### Limitations

This study has several limitations, the first of which was that it was done only with male runners and only at one speed. Hence, it is possible that the present results are limited to male runners of similar competitive levels running at a similar speed. It has been observed that lower limb strength levels and contact time may affect the optimal LBS between subjects ^33^. Further metatarsal phalangeal joint moments and optimal LBS are larger for heavier runners and at faster running speeds [for review: ^4^]. Our results should therefore be considered in relation to the population we studied (males with a body mass of 65.3 ± 5.7 kg) and for our testing speed of 13 km/h. In addition, the Control condition used was different from the two experimental conditions, although with similar mass and the same midsole material, but other characteristics such as geometry and midsole stiffness and compliance could affect the comparison.

## CONCLUSION AND PRACTICAL APPLICATIONS

Our results demonstrate that changes in LBS with curved carbon fiber plates in AFT shoes influence RE. This relationship suggests that moderately stiff (35.5 N/mm) shoes are most effective LBS to improve RE in AFT compared to stiffer (43.1 N/mm) and more flexible shoe conditions (20.1 N/mm). The improvements in the Stiff condition compared to Control condition seem not to be related to spatiotemporal changes (stride frequency and contact time), so future studies should perform more in-depth biomechanical analyses to identify differences running biomechanics with shoe with different LBS.

### Perspective

Most major running shoe companies now have a new style of road racing shoe with a thick lightweight midsole and a carbon fiber plate that increases the LBS of the shoe like the ones used in this study. Therefore, these results generate important new knowledge for the footwear industry regarding the possible optimal LBS to improve RE where intermediate stiffnesses were the most effective in improving RE. In addition, the improvement in RE was greater than reported in previous studies performed with flat plates or in traditional running shoes, so increasing LBS optimally in AFT has better results than in traditional running shoes.

## Acknowledgments

We thank the participants for participating. We thank *On running* for providing the shoes for this study without any conflict of interest or financial support.

